# Increasing cerebral blood flow improves cognition into late stages in Alzheimer’s disease mice

**DOI:** 10.1101/640912

**Authors:** Oliver Bracko, Brendah N. Njiru, Madisen Swallow, Muhammad Ali, Chris B. Schaffer

## Abstract

Alzheimer’s disease (AD) is associated with a 20-30% reduction in cerebral blood flow. In the APP/PS1 mouse model of AD, inhibiting neutrophil adhesion using an antibody against the neutrophil specific protein Ly6G was recently shown to drive rapid improvements in cerebral blood flow that was accompanied by an improvement in performance on short-term memory tasks. Here, in a longitudinal aging study, we assessed how far into disease development a single injection of anti-Ly6G can acutely improve memory function. We found that APP/PS1 mice as old as 15-16 months had improved performance on the object replacement and Y-maze tests of short-term memory, measured at one day after anti-Ly6G treatment. APP/PS1 mice 17-18 months of age or older did not show acute improvements in cognitive performance, although we did find that cerebral blood flow was still increased by 17% in 21-22 months old APP/PS1 mice given anti-Ly6G. These data add to the growing body of evidence suggesting that cerebral blood flow reductions are an important contributing factor to the cognitive dysfunction associated with neurodegenerative disease.

## Introduction

Alzheimer’s disease (AD) is the most common cause of dementia in the elderly, and is characterized by brain atrophy and a gradual cognitive decline that is correlated with a loss of synapses and death of neuronal cells, particularly in brain regions involved in learning and memory processes (Iadecola, 2017). Recent work has suggested that there are significant vascular contributions to the progression and severity of AD and other forms of dementia (Strickland, 2018). Cerebral blood flow (CBF) reductions ranging from 10–28% have been reported in patients with AD and other neurodegenerative diseases (Chen et al., 2011; Dopper et al., 2016; Hays et al., 2016; Leoni et al., 2017). These CBF reductions can be detected early in disease progression, before deposition of large amyloid plaques or significant neurological decline (Iturria-Medina et al., 2016; Kogure et al., 2000; Okonkwo et al., 2014). The degree of blood flow reduction correlates with the severity of some cognitive symptoms, suggesting that impaired CBF may directly contribute to cognitive decline in neurodegenerative disease. Decreased CBF in the thalamus and caudate was associated with poorer performance on a broad test of cognitive function in patients with AD and mild cognitive impairment (MCI) (Nation et al., 2013). In MCI patients, increased CBF was associated with better memory performance, while decreased CBF was correlated to poorer visuospatial perception and general cognitive dysfunction (Bangen et al., 2012; Bangen et al., 2017; Yoon et al., 2012). In addition, in aged humans without neurodegenerative disease, lower blood flow in hippocampus was associated with poorer spatial memory (Heo et al., 2010). Finally, reduced CBF was found to be predictive of later cognitive decline in healthy individuals (Xekardaki et al., 2015).

Mouse models of Alzheimer’s disease, which overexpress mutated forms of amyloid precursor protein (APP), similarly display CBF reductions of ~25% (Li et al., 2014; Ni et al., 2018; Niwa et al., 2002; Wiesmann et al., 2017), which occur early in the course of disease progression (Cruz Hernández et al., 2019; Li et al., 2014). In these mouse models of AD, increased CBF, achieved through increased physical activity, was correlated with enhanced memory function (Choi et al., 2018). In addition, in wild-type (WT) mice with chronic brain hypoperfusion show impaired memory function (Hattori et al., 2016; Wang et al., 2016). Taken together, these data suggest that overexpression of mutated forms of APP in mice leads to CBF deficits and such blood flow deficits contribute to cognitive dysfunction.

Several mechanisms have been proposed that may contribute to the CBF reductions seen in AD patients and mouse models, including tonic constriction of brain arterioles and reduced vascular density in the brain (Kisler et al., 2017). Vascular amyloidosis as well as oxidative stress and dysfunction in endothelial cells have been correlated with CBF reductions in AD (Hachinski et al., 1975; Iturria-Medina et al., 2016; Rogers et al., 1986), suggesting increased vascular inflammation may play a role in CBF reduction (Iadecola, 2017). Consistent with this idea, we recently found that most of the CBF reduction seen in mouse models of AD is caused by neutrophils that transiently adhere to the endothelial cell wall in brain capillaries. About 1.8% of cortical capillaries had stalled blood flow in the APP/PS1 mouse model of AD, as compared to 0.4% in WT mice. The incidence of capillary stalls could be reduced by treating with an antibody against the neutrophil-specific GPI-linked protein lymphocyte antigen 6 complex, locus G (Ly6G). This antibody interfered with neutrophil adhesion immediately, leading to a ~60% reduction of the number of capillary stalls and a resulting ~30% increase in CBF in APP/PS1 mice that occurred within minutes. Correlated to this rapid CBF recovery, we observed an improvement in spatial and working short-term memory within 3 hours in mice that were 10-11 months of age, shortly after cognitive deficits are first observed in this AD mouse model. Within a day the anti-Ly6G depleted neutrophils, which preserved the reduction in stalled capillaries, thereby maintaining the increased cerebral blood flow and improved memory performance (Cruz Hernández et al., 2019).

In this paper, we explore the correlation between CBF and cognition in APP/PS1 mice and aim to determine how far into disease progression treatment with anti-Ly6G can still lead to improvements in short-term memory as well as increased CBF.

## Materials and methods

### Animals

All animal procedures were approved by the Cornell Institutional Animal Care and Use Committee and were performed under the guidance of the Cornell Center for Animal Resources and Education (protocol number 2015-0029). We used aged transgenic mice (B6.Cg-Tg (APPswe, PSEN1dE9) 85Dbo/J, RRID: MMRRC_034832-JAX, The Jackson Laboratory) as a mouse model of AD (Jankowsky et al., 2004) and WT littermates (C57BL/6) as controls. Animals were of both sexes (Group 1 (APP/PS1): 4 females and 7 males; Group 2 (APP/PS1): 5 females and 6 males; and Group 3 (WT): 6 females and 6 males). One APP/PS1 mouse from Group 1 died at 14 months of age and one WT mouse from Group 3 died at 16 months of age.

### Behavior Experiments

We used two measures of short-term memory function, the object replacement task and the Y-maze task. All experiments were performed in an isolated room under red light conditions. Mice were habituated to the room where behavioral experiments were conducted over three visits to the room. During behavioral experiments, mice were brought to the behavior room one hour prior to starting the experiment to give animals time to adjust. Behavioral analysis was conducted at baseline and/or at 24 hours after intraperiotenial (i.p.) injection with α-Ly6G or isotype control antibodies (IP 4 mg/kg; α-Ly6G, clone 1A8, No. 561005; Iso-Ctr, Rat IgG2a, κ, No. 553929; BD Biosciences). Animals were 9-10 months of age at the start of the experiment.

The object replacement (OR) task evaluated spatial memory performance. All pairs of objects used for this experiment were tested previously in an independent cohort of mice to avoid any object or position bias (Cruz Hernández et al., 2019). Exploration time was defined as any time when there was physical contact with the object (whisking, sniffing, rearing on, or touching the object) or when the animal was oriented toward the object and the head was within 2 cm of the object. In trial 1, mice were allowed to explore two identical objects for 10 min in the arena and then returned to their home cage for 60 min. Mice were then returned to the testing arena for 3 min with one object moved to a novel location (trial 2). Care was taken to ensure that the change of placement altered both the intrinsic relationship between the objects (e.g. a rotation of the moved object) and the position of the moved object relative to visual cues (each wall of the testing arena had a distinct pattern). After each trial the arena and objects were cleaned using 70% ethanol. The preference score (%) for OR the task quantifies the degree of preference the mouse shows for the moved object and was calculated as ([exploration time of the moved object]/[exploration time of both objects]) × 100 from the data in trial 2.

The Y-Maze task was used to measure working memory by quantifying spontaneous alternation between arms of the maze. The Y-maze consisted of three arms at 120° and was made of light grey plastic. Each arm was 6-cm wide and 36-cm long and had 12.5-cm high walls. The maze was cleaned with 70% ethanol after each mouse. A mouse was placed in the Y-maze and allowed to explore for 6 min. A mouse was considered to have entered an arm if the whole body (except for the tail) entered the arm and to have exited if the whole body (except for the tail) exited the arm. If an animal consecutively entered three different arms, it was counted as an alternating trial. Because the maximum number of such alternating triads is the total number of arm entries minus 2, the spontaneous alternation score (%) was calculated as (the number of alternating triads)/(the total number of arm entries – 2) × 100.

Behavioral data for the OR and Y-maze tasks was quantified in two ways. First, the position of the nose and body of the mice was traced using Viewer III software (Biobserve, Bonn, Germany), and these positions were used together with the criteria described above to automatically score behavior. In addition, the videos were manually scored, again using the criteria described above, by a blinded experimenter. Automated tracking and manual scoring yielded similar results across all groups, so we report the automated tracking results.

### Surgical preparation and in vivo imaging of blood flow and vessel diameter

The surgery and in vivo two-photon microscopy were performed as previously described (Cruz Hernández et al., 2019). In brief, a 6-mm diameter craniotomy was prepared over parietal cortex and covered by gluing a glass coverslip to the skull. Animals rested for at least 3 weeks before imaging to minimize the impact of post-surgical inflammation. During the imaging session, mice were placed on a custom stereotactic frame and were anesthetized using ~1.5% isoflurane in 100% oxygen, with small adjustments to the isoflurane level made to maintain the respiratory rate at ~1 Hz. The mouse was kept at 37 °C with a feedback-controlled heating pad (40-90-8D DC, FHC). After being anesthetized and every hour during the experiment, mice received a sub-cutaneous injection of atropine (0.005 mg/100 g mouse weight; 54925-063-10, Med Pharmex) to prevent lung secretions and of 5% glucose in saline (1 mL/100 g mouse weight) to prevent dehydration. Texas Red dextran (40 μl, 2.5%, molecular weight = 70,000 kDa, Thermo Fisher Scientific) in saline was injected retro-orbitally just prior to imaging to label the vasculature. Mice were imaged on a locally-designed two-photon excited fluorescence (2PEF) microscope, using 900-nm, femtosecond duration laser pulses for excitation (Vision II, Coherent) and collecting Texas-Red emission through an interference filter with a center wavelength of 640 nm and a 75-nm bandwidth. We collected three-dimensional image stacks and identified penetrating arterioles based on their connectivity to readily-identifiable surface arterioles and by determining their flow direction using the motion of unlabeled red blood cells in the sea of fluorescently-labeled blood plasma. In each mouse, we identified 6-10 penetrating arterioles and took small three-dimensional image stacks to determine the diameter as well as line-scans along the centerline of the vessel to determine red blood cell flow speed (Santisakultarm et al., 2012). These measures were taken at baseline and ~1 hour after injection of α-Ly6G or isotype control antibodies (IP 4 mg/kg).

### Statistical Analysis

Behavioral data and measurements of changes in vessel diameter and blood flow were plotted using Tukey boxplots, where the box contains the middle two quartiles of the data, the whiskers extend 1.5 times the difference between the 75th and 25th percentiles, the black line defines the median, and the red line defines the mean, which is calculated excluding any outliers that sit outside the whiskers. Normality was tested with the D’Agostino-Pearson test and the data for each measurement of each group was found to be normally distributed. The statistical significance of differences between groups was evaluated by one-way analysis of variance (ANOVA) followed by pairwise comparisons using the multiple-comparison corrected Holm-Šídák test. To compare baseline blood flow (Fig. 4B), flow speed was fit to vessel diameter using linear regression and the models compared for APP/PS1 and WT mice. P-values <0.05 were considered statistically significant. All statistical analyses were performed using Prism7/8 software (GraphPad Software) or JMP Pro 14 (SAS Institute Inc.).

## Results

To evaluate how far into disease development increasing blood flow can improve cognitive symptoms in mouse models of Alzheimer’s disease, we treated APP/PS1 mice every two months with antibodies against Ly6G (α-Ly6G; 4 mg/kg animal weight, i.p.) or with isotype control antibodies (Iso-Ctr; 4 mg/kg, i.p.) and measured the impact on spatial and working memory. To control for the impact of repeated antibody treatment, the APP/PS1 mice were broken into two cohorts, each alternating between α-Ly6G and Iso-Ctr at successive treatments beginning with Group 1 (APP/PS1) receiving α-Ly6G and Group 2 (APP/PS1) receiving Iso-Ctr antibodies (Fig. 1). A third cohort mice, Group 3 (WT) received α-Ly6G at each treatment as a control. Animals began the study at 9-10 months of age, when short-term memory deficits first become evident in APP/PS1 mice (Serneels et al., 2009). For the first two time points (10-11 and 12-13 months of age), cognitive testing was performed both before and one day after antibody administration, while for the subsequent four time points cognitive testing was done only one day after antibody administration. Previous work by us (Cruz Hernández et al., 2019) and others (Daley et al., 2008) showed that circulating neutrophil counts are sharply reduced one day after treatment with this dose of α-Ly6G.

**Figure 1:**
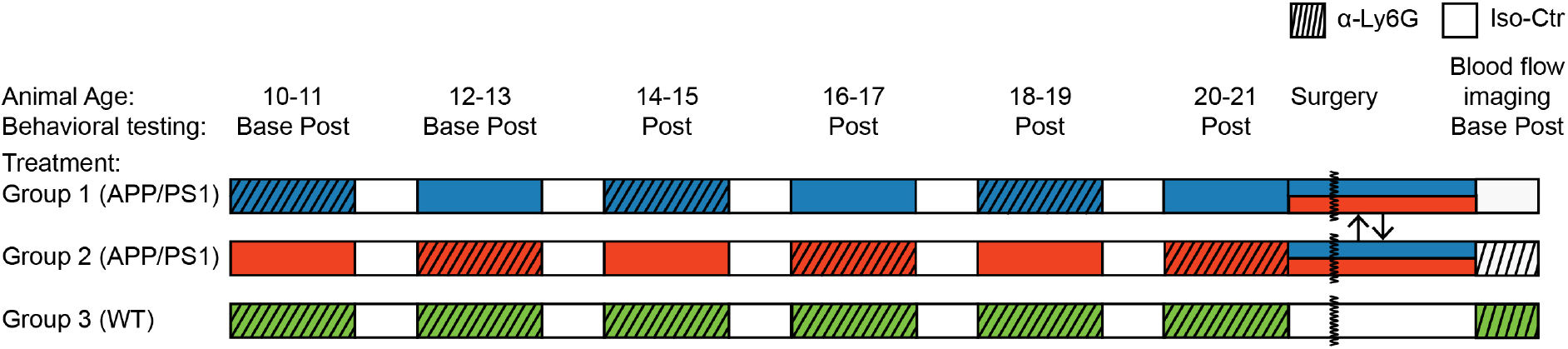
Experimental time line. Mice were divided into three groups: Group 1 (APP/PS1, blue), Group 2 (APP/PS1, red) and Group 3 (WT, green). At the first two timepoints (10-11 and 12-13 months of age) all groups of mice were tested using Y-maze and object replacement tests before and one day after treatment with α-Ly6G or Iso-Ctr antibodies. At subsequent time points, behavioral testing was done only one day after antibody treatment. Groups 1 (blue) and 2 (red) of APP/PS1 mice alternated between treatment with α-Ly6G and Iso-Ctr at each time point, as indicated, while Group 3 (green) of WT mice always received α-Ly6G. After the last behavioral testing at 20-21 months of age, the APP/PS1 mice were randomly regrouped and mice received a chronic craniotomy to enable measurement of blood flow speed in penetrating arterioles using 2PEF imaging. Blood flow was measured before and one-hour after treatment with α-Ly6G or Iso-Ctr antibodies in APP/PS1 mice, while WT mice all received α-Ly6G.

We used two tests of short-term memory, the Y-maze and the object replacement test. The object replacement test measures spatial memory by characterizing how much mice explore a novel placement of an object (Vogel-Ciernia and Wood, 2014). This is measured by first exposing mice to one spatial arrangement of two objects and then, after a 1-hour delay, quantifying the fraction of object exploration time mice spend on one object that has been moved to a novel location (Fig. 2A). For WT mice, about 60-75% of the time exploring the two objects will be spent on the moved object. The Y-maze measures working memory by characterizing how often mice explore areas where they have not been recently (Bak et al., 2017). This is measured by the percentage of entries into each arm of the three-arm maze that are part of an exploration that alternate between all three arms, called spontaneous alternation. For WT mice, 60-70% arm entries are part of such a spontaneous alternation triad.

**Figure 2:**
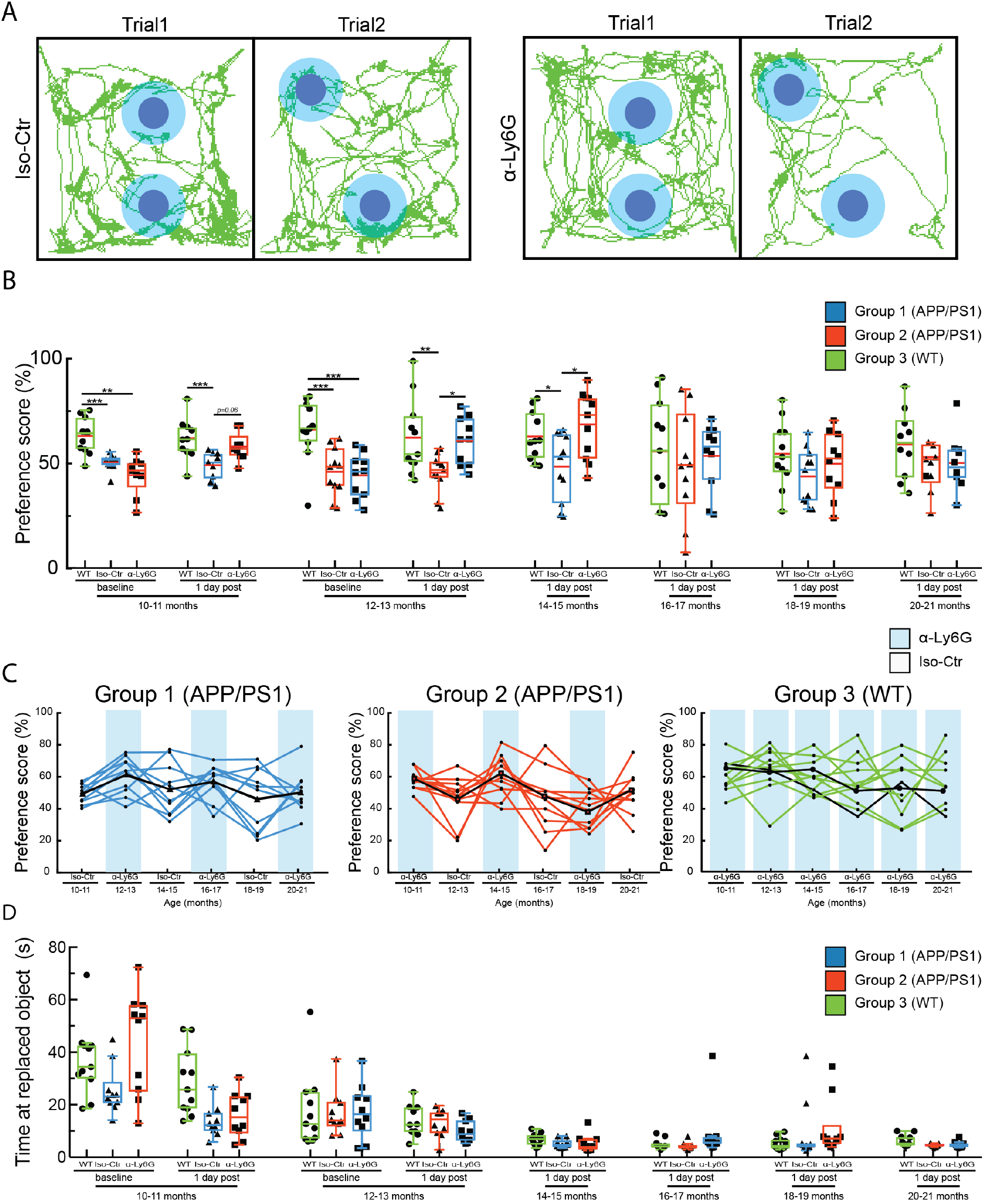
A single treatment with α-Ly6G restores spatial short-term memory one day after injection in APP/PS1 mice as old as 14-15 months of age. (A) Tracking of mouse nose location from video recording during training and trial phases of OR task taken one day after administration of α-Ly6G or Iso-Ctr antibodies in APP/PS1 mice at 14-15 months of age. (B) Preference score for OR of APP/PS1 and WT mice at baseline and at one day after a single administration of α-Ly6G or Iso-Ctr antibodies at different ages. (C) Preference score one day after treatment for individual animals from each group across age. Black line shows the median trend for each group. Blue shading indicates treatment at that time point with α-Ly6G while no shading indicates treatment with Iso-Ctr antibodies. (D) Time spent at the replaced object measured over 3 minutes for APP/PS1 and WT mice at baseline and at one day after a single administration of α-Ly6G or isotype control antibodies across animal age. (Group 1 (blue) APP/PS1: 10 mice; Group 2 (red) APP/PS1: 10 mice; Group 3 (green) WT: 11 mice; D’Agostino-Pearson was performed for normality followed by one-way ANOVA and Holms-Sidak’s multiple comparison; * p<0.05, ** p<0.01, *** p<0.001. All boxplots are defined as: whiskers extend 1.5 times the difference between the value of the 75th and 25th percentile, median=black line and mean= red line.)

In baseline measurements with 10-11- and 12-13-month old mice, we found a significant impairment of APP/PS1 mice (both Group 1 and Group 2) on both object replacement (Fig. 2B) and Y-maze tests (Fig. 3A), as compared to WT mice (Group 3). Notably, APP/PS1 mice from Group 1, which received α-Ly6G treatment at 10-11 months of age, showed deficits in both cognitive tests in the baseline measurement at 12-13 months of age, suggesting there was not a long-term cognitive impact of a single dose of α-Ly6G.

**Figure 3:**
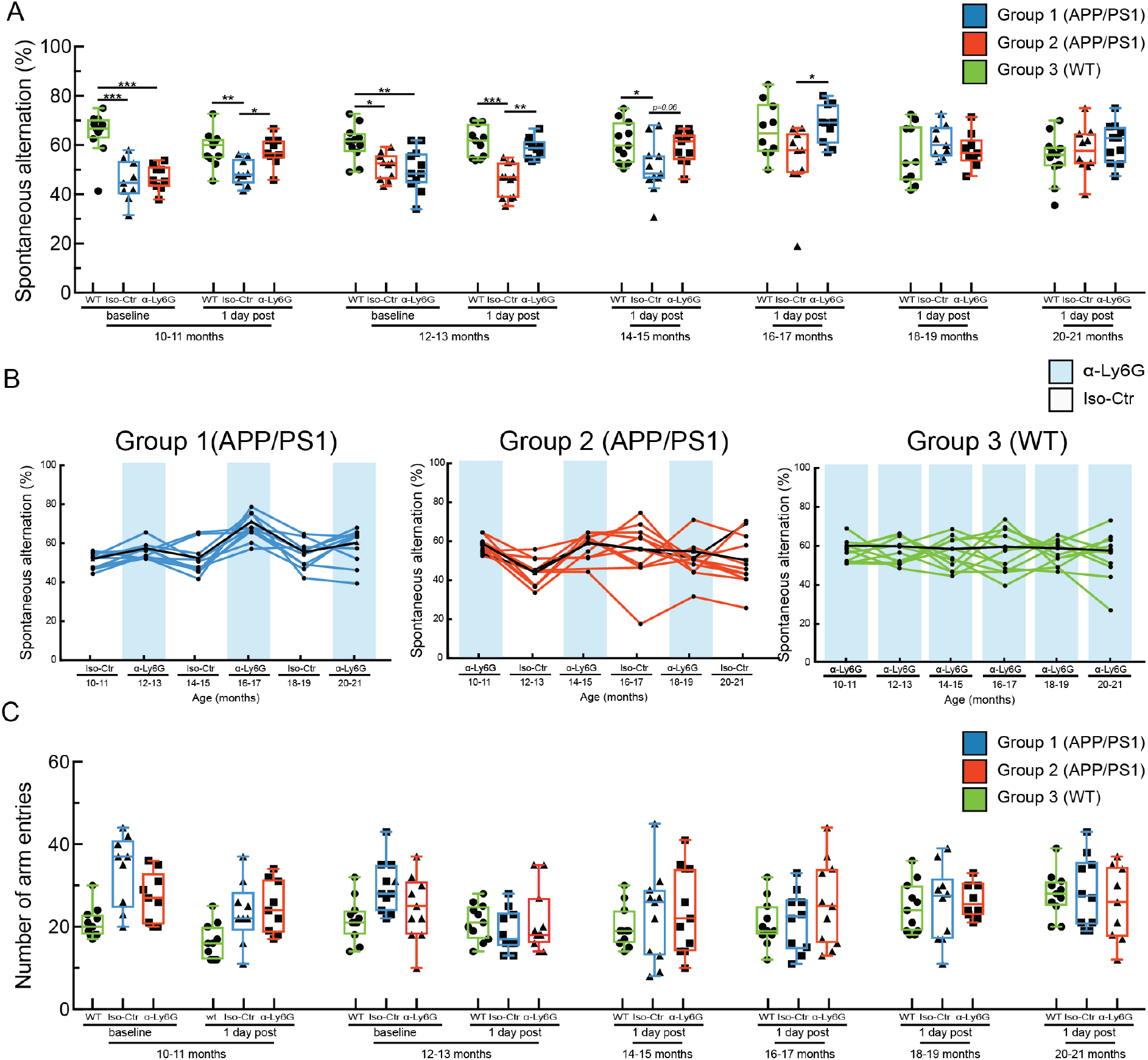
A single treatment with α-Ly6G restores working memory one day after injection in APP/PS1 mice as old as 16-17 months of age. (A) Spontaneous alternation in Y-maze task for APP/PS1 and WT mice at baseline and at one day after a single administration of α-Ly6G or Iso-Ctr antibodies at different ages. (B) Spontaneous alternation one day after treatment for individual animals from each group across age. Black line shows the median trend for each group. Blue shading indicates treatment at that time point with α-Ly6G while no shading indicates treatment with Iso-Ctr antibodies. (C) Number of arm entries for APP/PS1 and WT mice at baseline and at one day after a single administration of α-Ly6G or isotype control antibodies across animal age. (Group 1 (blue) APP/PS1: 10 mice; Group 2 (red) APP/PS1: 10 mice; Group 3 (green) WT: 11 mice; D’Agostino-Pearson was performed for normality followed by one-way ANOVA and Holms-Sidak’s multiple comparison; * p<0.05, ** p<0.01, *** p<0.001. All boxplots are defined as: whiskers extend 1.5 times the difference between the value of the 75th and 25th percentile, median=black line and mean= red line.)

APP/PS1 mice from both Groups 1 and 2 showed significantly improved cognitive performance one day after α-Ly6G treatment on the object replacement test at ages between 10-11 and 14-15 months of age (Fig. 2B and C) and on the Y-maze test at ages between 10-11 and 16-17 months of age (Fig. 3A and B), as compared to the corresponding Iso-Ctr treated APP/PS1 mice at each time point. In the object replacement task, there was no difference in α-Ly6G and Iso-Ctr treated APP/PS1 mice for animals older than 16-17 months and there was a trend toward decreased performance in WT animals, as well (Fig. 2B and C). In the Y-maze test, no difference between α-Ly6G and Iso-Ctr treated APP/PS 1 mice was observed for animals older than 18-19 months, and rather both groups of APP/PS1 showed slightly improved performance, irrespective of treatment. The performance of WT mice on the Y-maze test did not change with age (Fig. 3A and B). All mice showed decreased exploration time in the object replacement test with increasing age and more repetitions of the task (Fig. 2D), while all mice maintained steady levels of arm entries in the Y-maze task (Fig. 3C). Taken together, this suggests some saturation with this many repeats of the object replacement task (despite the use of new objects and object locations for each repetition), but no saturation of the Y-maze test. Nonetheless, there was still sufficient object exploration time in all repeats of the object replacement test to score the object preference.

In previous work, we used ASL-MRI to show that cortical perfusion was reduced by about 17% in APP/PS1 mice, as compared to WT controls, and that treatment with α-Ly6G in 6-8 month-old APP/PS1 mice led to a 13% increase in cortical perfusion, recovering about 2/3 of the blood flow deficit these mice exhibited relative to WT controls (Cruz Hernández et al., 2019). Using optical approaches to measure blood flow speed in individual cortical vessels, we also showed that the median blood flow speed in penetrating arterioles increased by ~30% in 3-4 and 11-14 month old APP/PS1 mice within an hour of treatment with α-Ly6G (Cruz Hernández et al., 2019). This increase in cerebral blood flow was correlated with a rapid improvement in short-term memory in APP/PS1 mice after α-Ly6G treatment. We wanted to understand if the loss of the rapid rescue of cognitive function after α-Ly6G treatment in older APP/PS1 shown here was a result of α-Ly6G treatment no longer leading to improved cerebral blood flow. We implanted chronic cranial windows and used two-photon excited fluorescence microscopy to measure blood flow speed in individual penetrating arterioles in all groups of mice when they reached 20-21 months of age (Fig. 4A). We pooled the APP/PS1 mice from Groups 1 and 2, identified 11 animals with high-quality craniotomies, split them into two groups (α-Ly6G, 6 mice; Iso-Ctr, 5 mice), and took measurements before and ~1 hour after treatment with α-Ly6G or Iso-Ctr antibodies. Five WT mice from Group 3 were, similarly, measured before and ~1 hour after α-Ly6G treatment. There was a trend showing that baseline blood flow speeds were about 14% lower, on average, in penetrating arterioles from aged APP/PS1 as compared to aged WT mice (Fig. 4b; p = 0.15, least squares fit of speed vs. diameter for APP/PS1 and WT mice). Treatment with α-Ly6G or isotype control antibodies did not, on average, change the diameter of penetrating arterioles in APP/PS1 or WT mice (Fig. 4C). There was a trend toward increased blood flow speed (Fig. 4D; p=0.07, one-way ANOVA with Holms-Sidak’s multiple comparison correction) and a significant increase of ~17% in the median volumetric blood flow in individual penetrating arterioles (Fig. 4E; p=0.04, one-way ANOVA with Holms-Sidak’s multiple comparison correction) in APP/PS1 mice treated with α-Ly6G, but not in WT mice treated with α-Ly6G or in APP/PS1 mice receiving isotype control antibodies.

**Figure 4:**
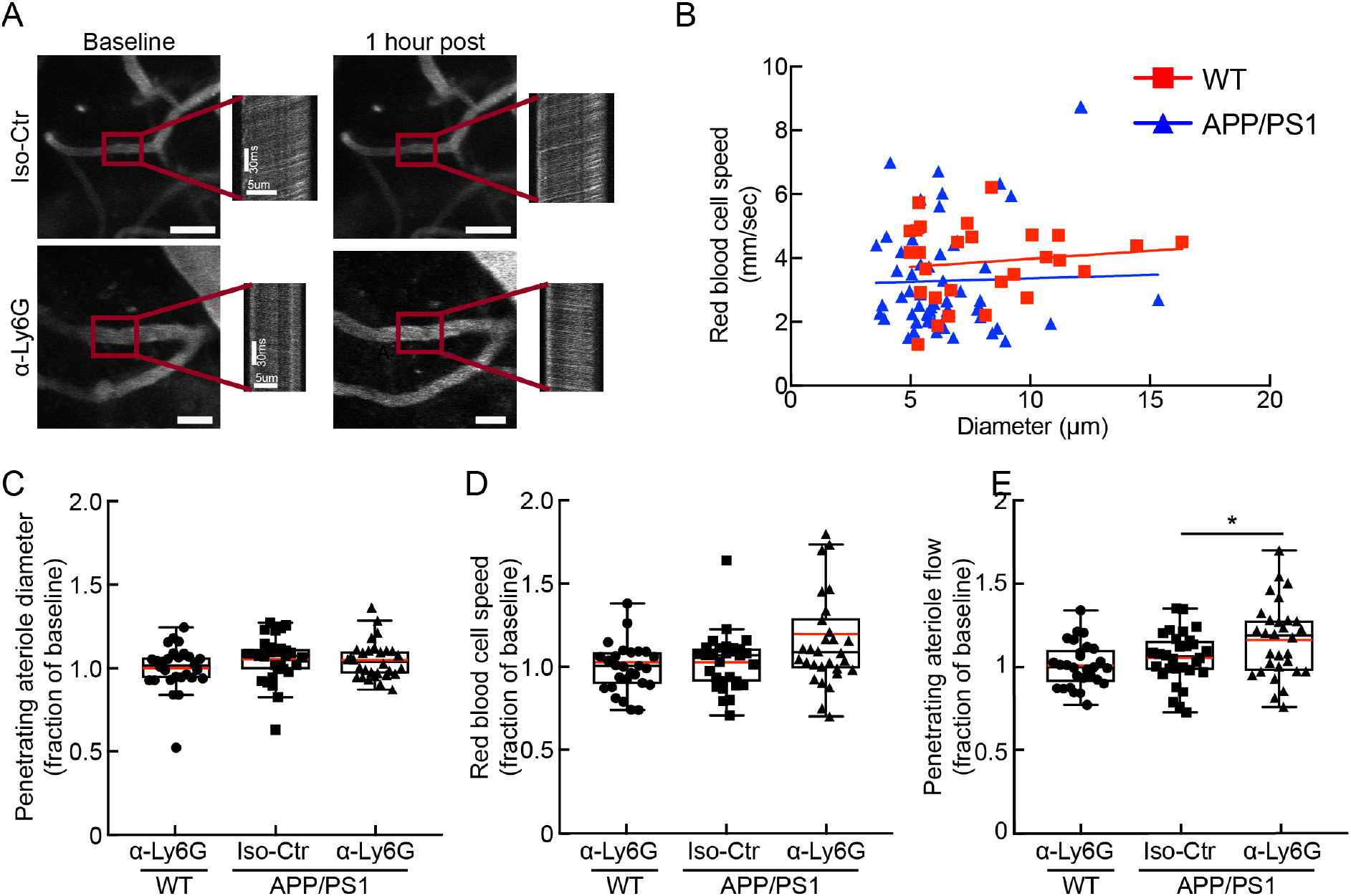
Administration of antibodies against Ly6G increased CBF in APP/PS1 mice at 21-22 months of age. (A) 2PEF images of cortical surface blood vessels and line scan data from penetrating arterioles (indicated with red box) from APP/PS1 mice taken at baseline and one hour after treatment with α-Ly6G or Iso-Ctr antibodies. Scale bar in images: 20μm. Scale bars are indicated in baseline line scan images and post treatment line scans are on same scale. (B) Scatter plot of baseline blood flow speed vs. vessel diameter for penetrating arterioles from 21-22-month-old WT mice (red) and 21-22-month-old APP/PS1 mice (blue). (APP/PS1: 55 arterioles across 11 mice; WT: 28 arterioles across 5 mice) (C) Vessel diameter, (D) RBC flow speed, and (E) Volumetric blood flow from penetrating arterioles after α-Ly6G or isotype control antibody administration in 20-21 months old APP/PS1 mice and WT control animals, shown as a fraction of baseline. (APP/PS1 α-Ly6G: 34 arterioles across 6 mice; APP/PS1 Iso-Ctr: 21 arterioles across 5 mice; WT α-Ly6G: 28 arterioles across 5 mice; D’Agostino-Pearson was performed for normality followed by one-way ANOVA and Holms-Sidak’s multiple comparison; * p<0.05. All boxplots are defined as: whiskers extend 1.5 times the difference between the value of the 75th and 25th percentile, median=black line and mean= red line.)

## Discussion

This study aimed to identify the stages of disease progression in which short-term memory was acutely improved after a single treatment with α-Ly6G in APP/PS1 mice. In previous work we showed that there are an increased number of transiently stalled capillaries in the cortex of APP/PS1 mice from a few months to two years of age, as compared to WT mice, and that these stalls were caused by neutrophils adhered to the capillary endothelium. In APP/PS1 mice across a broad range of ages from 3 to 15 months, we further found that administering antibodies against the neutrophil surface protein Ly6G led to the capillary stalls resolving and cerebral blood flow increasing within minutes. In mice that were 10-11 months of age, when short-term memory deficits first become evident in the APP/PS1 mice, a single treatment with α-Ly6G improved performance on spatial and working memory tasks within hours and this improvement persisted for at least one day (Cruz Hernández et al., 2019). Here, we extend this work in two important directions. First, we found that treatment with α-Ly6G still improved cortical blood flow in APP/PS1 mice even at an extreme age of 21-22 months. Taken together with our previous results, this data suggests that CBF deficits in this AD mouse model are dominated by capillary stalling throughout the life of the mouse. Second, we demonstrated that administration of α-Ly6G led to improved short-term memory function one day after treatment in APP/PS1 mice as old as 15-16 months of age, but not in older animals.

The rapid improvement in short-term memory performance in APP/PS1 mice after increasing CBF with α-Ly6G treatment suggests that neuronal networks in the brain of these animals are still largely intact but have impaired function due to decreased CBF. Improving CBF in APP/PS1 mice could acutely impact the function of neurons and neuronal networks in several ways. The ~20% CBF deficit APP/PS1 mice show relative to WT mice could impact neural function directly by interfering with normal metabolic homeostasis. Indeed, APP/PS1 mice have been shown to have deficits in glucose metabolism, mitochondrial function, and in ATP/ADP level regulation in astrocytes and neurons (Kang et al., 2017; Pathak et al., 2013; Venkat et al., 2016), which could all impair neural function. Decreased CBF could also cause mild hypoxia, which has been shown to selectively suppress the activity of inhibitory neurons, leading to hyperactivity in hippocampal networks that is associated with impaired cognitive performance (Haberman et al., 2017; Miller et al., 2008; Zhao and Gong, 2015). Supporting this idea, hyperactivity and epileptiform-like neural activity have been observed in APP/PS1 mice (Busche et al., 2019). Impaired CBF may also interfere with the synaptic plasticity necessary for learning and memory, and recent work has shown that synaptic plasticity is impaired as early as 6 months of age in APP/PS1 mice (Viana da Silva et al., 2016). The acute improvement in cognitive performance after α-Ly6G we observed is likely due to increased CBF rescuing some of these effects. However, treatment with α-Ly6G may have additional effects that influence brain function beyond increasing CBF, such as decreasing the infiltration of neutrophils into the brain and thus decreasing inflammation that may impact neural function (Zenaro et al., 2015).

The fact that a single treatment with α-Ly6G still improved CBF in APP/PS1 mice as old as 21-22 months of age, but short-term memory was not improved in mice older than 15-16 months of age suggests that other mechanisms contributing to cognitive dysfunction become more prominent at advanced stages of disease development. For example, structural abnormalities, such as dystrophic neurites, and the death of neurons and other brain cells are first observed in APP/PS1 mice at around 17 months of age (Blanchard et al., 2003; Clarke et al., 2018; Kitazawa et al., 2012; Wirths and Bayer, 2010), effects which an acute improvement in CBF would not be expected to rescue. Our data also does not rule out the possibility that improving cerebral blood flow over a longer period of time by, for example, repeat injections of α-Ly6G could lead to improvements in cognitive function in APP/PS1 older than 15-16 months of age.

Crucial remaining questions include whether the same mechanisms discussed here contribute to CBF reductions in AD patients and whether CBF increases could improve cognitive function in AD patients, at least in earlier stages of disease progression. In AD mouse models, we have shown that nearly all of the CBF deficit is due to leukocytes adhering to endothelial cells in brain capillaries, suggesting that vascular inflammation, induced by the toxic effects of amyloid-beta overexpression, is the ultimate cause. Chronic neuroinflammation is a well-recognized feature of AD in humans. In addition, recent work has specifically documented increased expression of inflammatory factors, including ICAM-1, VCAM-1, and TNF-α in brain endothelial cells from aged and AD patients (Grammas and Ovase, 2001; Smyth et al., 2018). Thus, the vascular inflammation that could lead to leukocyte adhesion and stalled blood flow in capillaries is present in AD patients. In the retina, a parallel story has already been established, where vascular inflammation secondary to diabetic retinopathy causes leukocyte adherence in retinal vessels that decreases retinal blood flow (Bek and Ledet, 1996; Burgansky-Eliash et al., 2010; Muir et al., 2012). If the blood flow deficits in AD patients are similarly caused by leukocyte adhesion in brain microvessels, therapeutic approaches that may decrease vascular inflammation or inhibit leukocyte adhesion should be pursued. As to the question of whether increasing CBF will improve cognition in AD patients, data from the last decade correlating slower cognitive decline with higher CBF suggests there is reason for hope (Bangen et al., 2017; Yoon et al., 2012). In support of this, limited studies in AD patients where a piece of omentum was surgically placing on the brain surface led to increased brain blood flow that was correlated to improvements in memory function (Goldsmith, 2017).

## Acknowledgements

We thank Nancy E. Ruiz-Uribe and Laibaik Park for critically reading this manuscript.

